# The proficiency of the original host species determines community-level plasmid dynamics

**DOI:** 10.1101/2020.10.13.337451

**Authors:** Anastasia Kottara, James P.J. Hall, Michael A. Brockhurst

## Abstract

Plasmids are common in natural bacterial communities, facilitating bacterial evolution via horizontal gene transfer. Bacterial species vary in their proficiency to host plasmids: Whereas plasmids are stably maintained in some species regardless of selection for plasmid-encoded genes, in other species, even beneficial plasmids are rapidly lost. It is, however, unclear how this variation in host proficiency affects plasmid persistence in communities. Here, we test this using multispecies bacterial soil communities comprising species varying in their proficiency to host a large conjugative mercury resistance plasmid. Plasmids reached higher community-level abundance where beneficial and when introduced to the community in a more proficient host species. Proficient plasmid host species were also better able to disseminate the plasmid to a wider diversity of host species. These findings suggest that the dynamics of plasmids in natural bacterial communities depend not only upon the plasmid’s attributes and the selective environment, but also upon the proficiency of their host species.

## INTRODUCTION

Mobile genetic elements (MGEs) like plasmids, temperate bacteriophages, and transposons, are important agents of horizontal gene transfer (HGT) driving the diversification of bacterial genomes (Frost *et al.* 2005; Hall, Brockhurst and Harrison 2017a; Brockhurst *et al.* 2019). Conjugative plasmids contain genes encoding core plasmid functions – including their own propagation, replication, stability and transfer – along with accessory genes that encode traits like antibiotic and metal resistance (Norman, Hansen and Sørensen 2009). While the plasmid’s accessory genes can directly benefit the host cell by providing them with new ecological functions, the plasmid’s core functions can impose a heavy burden on the host cell, the accessory genes can directly benefit the host cell by providing them with new ecological functions (Baltrus 2013; San Millan and Maclean 2017). Mathematical models of plasmid population dynamics suggest that the plasmid cost, conjugation rate, segregation rate, and the strength of positive selection are key parameters determining plasmid survival in bacterial populations (Stewart and Levin 1977; Levin, Stewart and Rice 1979; Simonsen *et al.* 1990; Bergstrom, Lipsitch and Levin 2000).

Plasmids are expected to spread under positive selection for their encoded accessory genes (San Millan *et al.* 2014; Harrison *et al.* 2015), however, because accessory genes can be captured by the bacterial chromosome rendering the plasmid redundant, positive selection does not guarantee the long-term survival of plasmids (Bergstrom, Lipsitch and Levin 2000). Meanwhile, in the absence of positive selection, plasmids are expected to go extinct due to purifying selection because the benefits of accessory genes do not outweigh the costs of plasmid carriage (Bergstrom, Lipsitch and Levin 2000). Since rates of conjugation appear to often be too low for plasmids to persist as infectious elements (although see: Lopatkin *et al.* (2017) and Stevenson *et al.* (2017)), it has been argued that the widespread distribution of plasmids is paradoxical (the plasmid paradox: Harrison and Brockhurst (2012)). Yet, plasmids have been found to stably persist in natural bacterial communities in the absence of measurable positive selection, where the factors allowing plasmid stability are puzzling (Heuer and Smalla 2012).

Most studies of plasmid dynamics focus on a single-host species, whereas, in natural bacterial communities, many potential host species co-exist, potentially broadening the range of conditions under which plasmids can survive. This limitation of current understanding is particularly interesting considering that several studies have shown that plasmids are not equally stable across host species (De Gelder *et al.* 2007; Kottara *et al.* 2018; Sakuda *et al.* 2018). For example, while the mercury resistance plasmid pQBR103 was highly stable for >400 generations with or without mercury selection in *P. fluorescens* and *P. savastanoi*, it was unstable to varying degrees in *P. stutzeri* (generally lost within ~100-400 generations), *P. aeruginosa* and *P. putida* (<6 generations) even with strong mercury selection (Kottara *et al.* 2018).

Hall *et al.* (2016) showed, by tracking the dynamics of the mercury resistance plasmid pQBR57 in a two-species soil community of *P. fluorescens* and *P. putida*, that between-species transfer of the plasmid from a proficient host, *P. fluorescens*, to an unstable host, *P. putida*, allowed the plasmid to persist in *P. putida* both with and without mercury selection. This finding suggests that the dynamics of a plasmid in a bacterial community is likely to depend on the proficiency of the plasmid host species to stably maintain the plasmid. This leads to the prediction that, at the community-level, plasmid abundance will be higher in communities where it is carried by a proficient original plasmid host, since this species will both be able to maintain the plasmid in its own population, and then disseminate the plasmid to other species in the community.

To test this prediction, we tracked the dynamics of pQBR103 in a three-species community of *P. fluorescens*, *P. stutzeri* and *P. putida* with and without mercury selection. We varied which of the species carried the plasmid at the start of the experiment. We hypothesised that the community-level plasmid abundance would vary according to the proficiency of the original plasmid host species to act as hosts to pQBR103, which varies hierarchically – *P. fluorescens* > *P. stutzeri* > *P. putida* (Kottara *et al.* 2018). Replicate communities were propagated in effectively sterile potting soil microcosms, which provide spatial structure and a low resource environment that more closely resemble the natural physical and chemical conditions in soil and promote the stable co-existence of multiple bacterial species (Gómez and Buckling 2011; Heuer and Smalla 2012; Hall *et al.* 2015; Hall *et al.* 2016).

## MATERIALS AND METHODS

### Bacterial strains and plasmid

Three *Pseudomonas* species – *P. fluorescens* SBW25 (Rainey, Bailey and Thompson 1994), *P. stutzeri* JM300 (DSM 10701) (Busquets *et al.* 2012) and *P. putida* KT2440 (Bagdasarian *et al.* 1981) – were utilised in this study. *Pseudomonas* species were labelled by directed insertion of either streptomycin (Sm^R^) or gentamicin resistance (Gm^R^) marker using the mini-Tn*7* transposon system (Lambertsen, Sternberg and Molin 2004). The plasmid used in this study, pQBR103 is a large conjugative plasmid (425 kb) that confers mercury resistance via a *mer* operon encoded on a Tn*5042* transposon (Lilley *et al.* 1996; Tett *et al.* 2007). pQBR103 plasmid is part of a group of 136 plasmids that were isolated from the bacterial community inhabiting the sugar beet rhizosphere and phyllosphere during a long-term field experiment (Lilley *et al.* 1996). pQBR103 was acquired by conjugation into labelled strain of *P. fluorescens* that was introduced onto the naturally occurring bacterial community colonising the sugar beet rhizosphere with the primary plasmid-host remaining unknown (Lilley *et al.* 1996). To obtain the initial plasmid-bearing clones of each host species to start the selection experiment, pQBR103 plasmid was conjugated into *P. stutzeri* Gm^R^, *P. putida* Sm^R^ and *P. fluorescens* Sm^R^*lacZ* from the plasmid-bearing *P. fluorescens* SBW25 Sm^R^ or Gm^R^ stocks. Plasmid conjugation was performed by mixing 1:1 each of the plasmid-free with the plasmid-bearing strains, incubating for 48 h and spreading on King’s B growth (KB) agar plates containing 5 μg mL^−1^ gentamicin or 50 μg mL^−1^ streptomycin (50 μg mL^−1^ X-Gal) and 20 μM of mercury(II) chloride to select for transconjugant colonies (Simonsen *et al.* 1990). The conjugation assays were conducted in 6 mL KB medium in 30 mL universal vials (‘microcosms’) at 28°C in shaking conditions (180 rpm).

### Selection experiment

To account for the high segregation rate of the plasmid in *P. putida* KT2440 (Kottara *et al.* 2018) and to ensure high starting frequencies of plasmid carriage across all the tested bacterial strains, single colonies of each plasmid-bearing species were reconditioned overnight and then transferred in fresh media containing mercury. Specifically, individual colonies (n=12) of each plasmid-bearing *Pseudomonas* species were picked into separate 6 mL KB microcosms and incubated overnight at 28°C with shaking 180 rpm after which time 1% of each population was transferred to grow for 24 h in fresh KB microcosms containing 50 μM of mercury(II) chloride at same temperature and shaking conditions; this concentration of mercury was used to select for the pQBR103 plasmid based on previous findings (Kottara *et al.* 2018). Similarly, 24 colonies of each plasmid-free *Pseudomonas* species were each grown overnight in KB 6 mL microcosms and transferred to grow for 24 h in fresh KB microcosms at same temperature and shaking conditions.

#### Bacterial communities

We used soil microcosms to evolve three different bacterial communities differing by which species carried the plasmid at the beginning of the experiment (original plasmid host). To prepare the soil microcosms, we added 10 g of John Innes No. 2 compost soil in 30 mL universal vials which we autoclaved twice. By autoclaving the compost soil two times, we established an effectively sterile micro-environment with the physical and chemical properties of soil which did not contain other culturable bacteria than our inoculum (Gómez and Buckling 2010; Hall *et al.* 2015; Hall *et al.* 2016). Three different bacterial communities were then constructed: *P. fluorescens* (carrying pQBR103) with *P. stutzeri* and *P. putida*; *P. fluorescens* with *P. stutzeri* (pQBR103) and *P. putida*; *P. fluorescens* with *P. stutzeri* and *P. putida* (pQBR103). Six replicates of each community were grown either without mercury or with mercury (16 μg g^−1^ Hg(II)); this concentration of mercury was used to select for the pQBR103 plasmid while could allow the survival of the plasmid-free species based on previous findings (Hall *et al.* 2015). Each community had a starting ratio of 1:1:1 of each *Pseudomonas* species such that the starting frequency of pQBR103 in the community was approximately 33%. To remove spent media and residual mercury from overnight cultures each inoculum was briefly vortexed, then centrifuged for 1 min at 10,000 rpm and resuspended in 1 mL M9 salt solution (Cold Spring Harbor Protocols). 100 μL was then inoculated into soil microcosms and incubated at 28°C at 80% humidity (Hall *et al.* 2016).

#### Serial transfers and bacterial counts

Every 4 days, 10 mL of M9 buffer and 20 glass beads were added to each soil microcosm and mixed by vortexing for 1 min, and 100 μL of soil wash was transferred to a fresh soil microcosm as previously described by Hall *et al.* (2016). Bacterial counts for each species were estimated by plating onto selective media: 50 μg mL^−1^ streptomycin + 50 μg mL^−1^ X-Gal KB agar plates and 5 μg mL^−1^ gentamicin KB agar plates, each of which was then replica plated onto mercury KB agar plates (100 μM mercury(II) chloride). The bacterial communities were evolved for 10 transfers (~40 days).

#### Plasmid and mercury-transposon screening

Twenty-four mercury-resistant colonies of each *Pseudomonas* species were sampled every 2 transfers from the mercury containing plates and tested for the presence of the plasmid and mercury transposon by PCR screening. The PCR used the same sets of primers as previously described [*mer* operon on the Tn*5042* transposon – forward primer: TGCAAGACACCCCCTATTGGAC, reverse primer: TTCGGCGACCAGCTTGATGAAC and origin of replication of the plasmid (*ori*V) – forward primer: TGCCTAATCGTGTGTAATGTC, reverse primer: ACTCTGGCCTGCAAGTTTC] (Harrison *et al.* 2015; Kottara *et al.* 2018).

### Statistics

Statistical analyses were performed using RStudio version 3.2.3 (R Core Team 2013). Shapiro-Wilk test, normal Q-Q plots, histograms and box-plots were used to examine the normality of the data. We found that in most cases the data were not normally distributed, and in such cases used a non-parametric test. Cumulative plasmid abundance in each community over time was estimated as the area under the curve using the function *auc* of the package ‘flux’ (Jurasinski, Koebsch and Hagemann 2012). Community-level plasmid abundances in the plasmid host treatments were compared by using the Kruskal-Wallis test. To assess the plasmid-dynamics within each species, we compared plasmid frequencies in the plasmid-recipient species population as the area under the curve. The integral estimates of the plasmid frequency in the recipient species were compared between the mercury conditions using the Kruskal-Wallis test. To assess the timing of chromosomal acquisition of the mercury transposon Tn*5042* in *P. putida* differed between the plasmid host treatments, for each population we recorded the transfer number when we first observed plasmid-free transposon-containing genotypes of *P. putida*. We compared these values between the plasmid host treatments using the Kruskal-Wallis test. The species diversity of plasmid-carriers was calculated as the 1 - D Simpson’s Index, 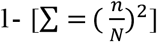 where, n = the end-point population density of each plasmid-bearer species in community, and, N = the end-point population density of all plasmid-bearer species. We compared diversities between the plasmid host treatments and mercury conditions by using the Kruskal-Wallis test.

## RESULTS

### Original plasmid host species identity affects community-level plasmid abundance

The bacterial host species vary in their ability to stably maintain pQBR103 hierarchically as follows: *P. fluorescens* > *P. stutzeri* > *P. putida* (Kottara *et al.* 2018). We hypothesised therefore that the identity of the original plasmid host in a community is likely to affect the dynamics of the plasmid-encoded mercury resistance at the community-level. To test this, we quantified the total plasmid abundance in each community (Figure 1). Mercury selection increased total plasmid abundance (effect of mercury; 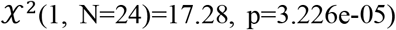 and total plasmid abundance varied with original plasmid host identity, such that both with (effect of plasmid treatment; 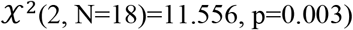 and without (effect of plasmid treatment; 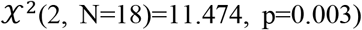 mercury selection, the total plasmid abundance was higher when the original plasmid host was *P. fluorescens*. Together these data suggest that community-level plasmid dynamics are affected by both the positive selection for plasmid-encoded traits and the identity of the original plasmid host species, being enhanced when plasmids are beneficial and carried by a proficient plasmid host.

**Figure 1.**
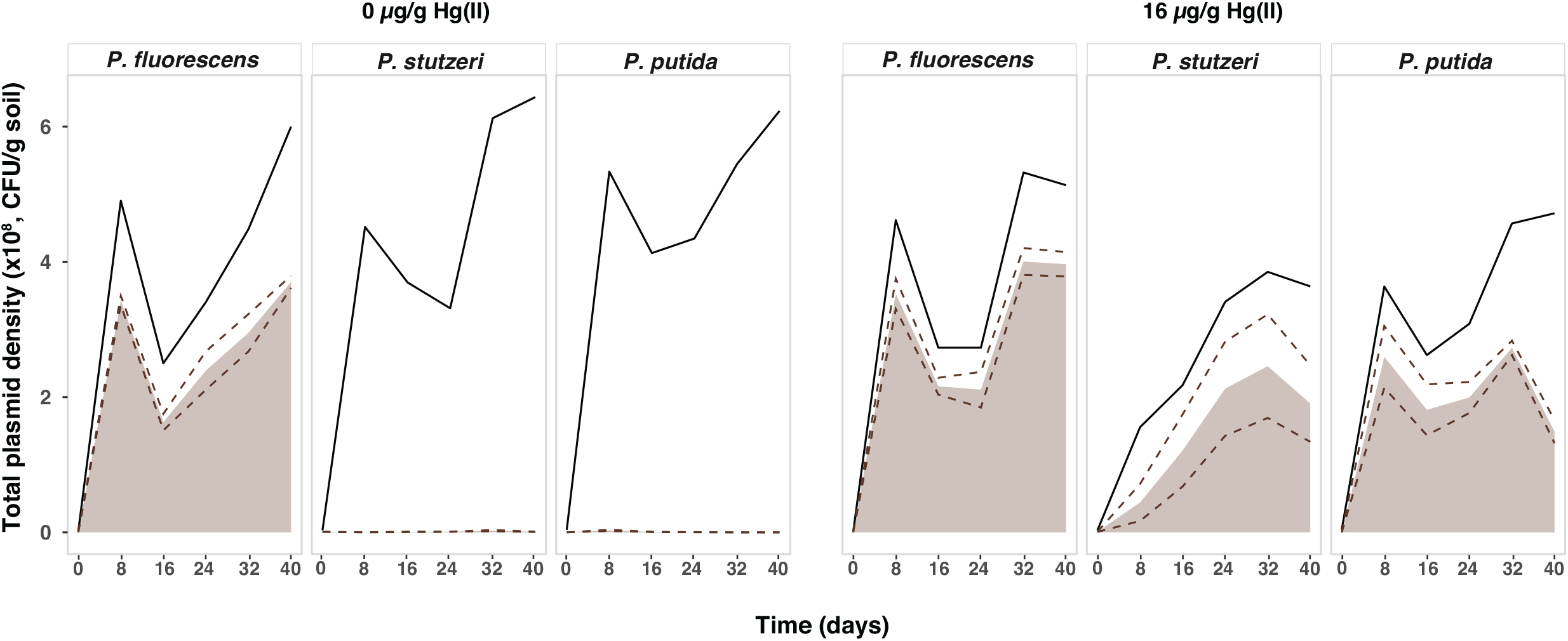
Total plasmid density in the community throughout the selection experiment. Panels data for communities that varied in mercury selection (without mercury, left-hand set; with mercury, right-hand set) their initial original plasmid host (from left to right in each set: *P. fluorescens*, *P. stutzeri* or *P. putida*). Brown shaded area shows the mean plasmid abundance in the community ± standard error (dotted line) from six replicates. Solid lines show the mean total community bacterial density from six replicates.

### Species-level plasmid dynamics within communities

To understand how the variation in community-level plasmid abundance was driven by original plasmid host identity, we next examined the species-level plasmid dynamics in each community. As predicted, when a proficient plasmid-host – *P. fluorescens* – was the original plasmid host it maintained the plasmid at high frequency within its population both with and without mercury (Figure 2). We detected plasmid dissemination from *P. fluorescens* to the other species at higher frequencies under mercury selection (effect of mercury; 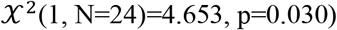. This occurred to *P. putida* in all replicates and to *P. stutzeri* in 2/6 replicates with mercury selection and also to *P. stutzeri* at low levels in some of the communities without mercury selection. When *P. stutzeri* was the original plasmid host, it also maintained the plasmid within its own population both with and without mercury, and disseminated plasmids to the other species at a higher rate with mercury (effect of mercury; 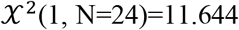 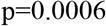 (Figure 3). Variation in total plasmid abundance between replicate communities in this treatment appear to have been caused by whether or not *P. fluorescens* acquired the plasmid before it was driven extinct by toxic mercury: where transmission to *P. fluorescens* occurred, total plasmid abundances were higher (Figure 3). Where *P. putida* was the original plasmid host, it did not maintain the plasmid within its own population: without mercury, the plasmid was simply lost, whereas, with mercury, plasmid-bearers were replaced by mutants that had inserted the Tn*5042* encoding the *mer* operon into their chromosome (Figure 4). Chromosomal insertions of the Tn*5042* in *P. putida* were observed in the other plasmid host treatments, but arose much later in these communities where *P. putida* had to acquire the plasmid horizontally from either *P. fluorescens* or *P. stuzeri* (effect of treatment; 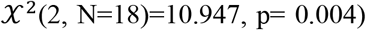. Although *P. putida* eventually lost the plasmid from its own population, prior to this loss it successfully disseminated the plasmid to *P. fluorescens* in 6/6 replicates and to *P. stutzeri* in 3/6 replicates with mercury selection (Figure 4).

**Figure 2.**
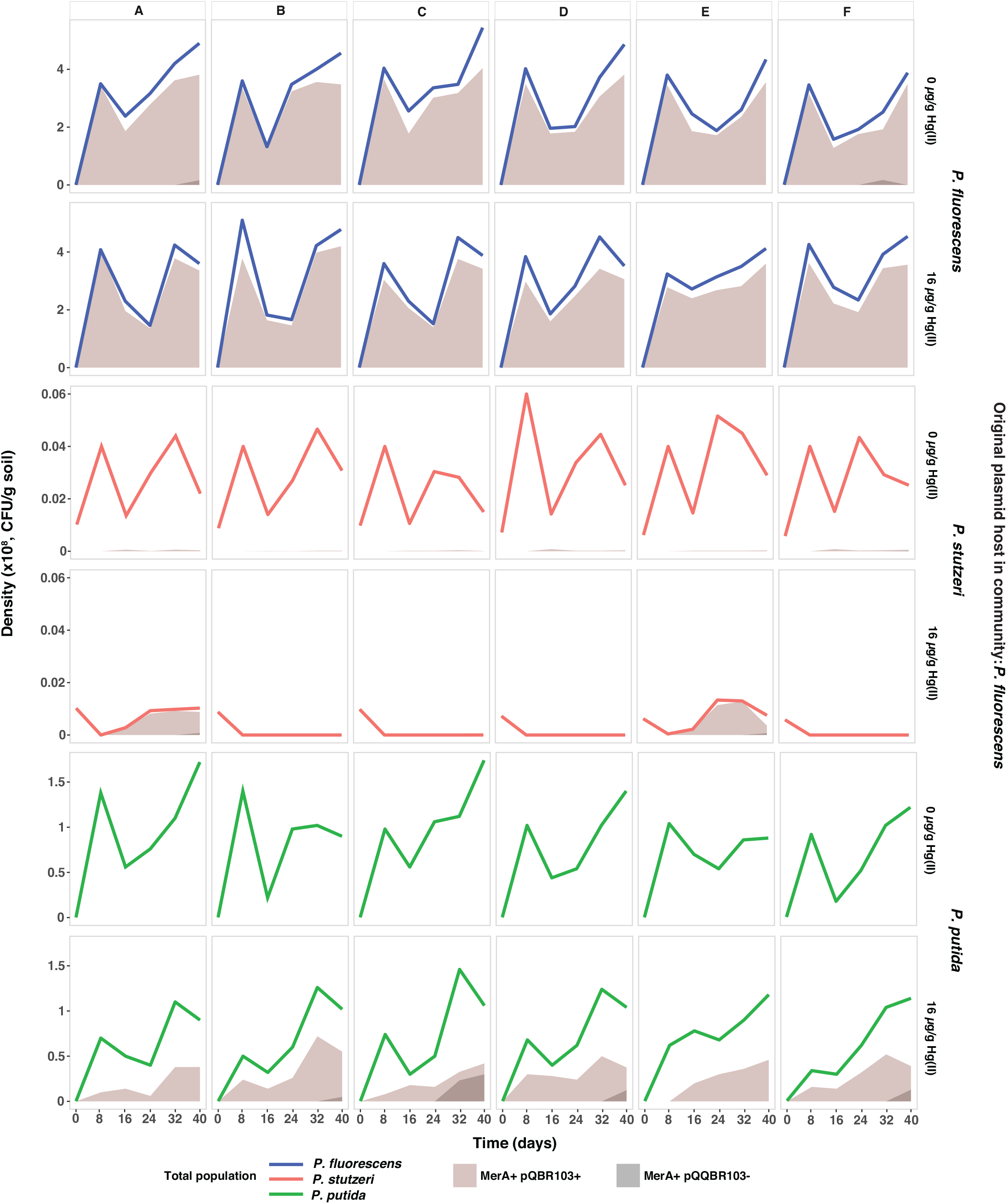
Population density and mobile genetic element dynamics in communities where *P. fluorescens* was the original plasmid host. A-F clonal populations evolving with or without mercury. Lines show the population densities of *P. fluorescens* (blue); *P. stutzeri* (red); *P. putida* (green). Brown areas show the density of plasmid carriers; Grey areas show the density of cells that have retained the Tn5042 but lost the plasmid.

**Figure 3.**
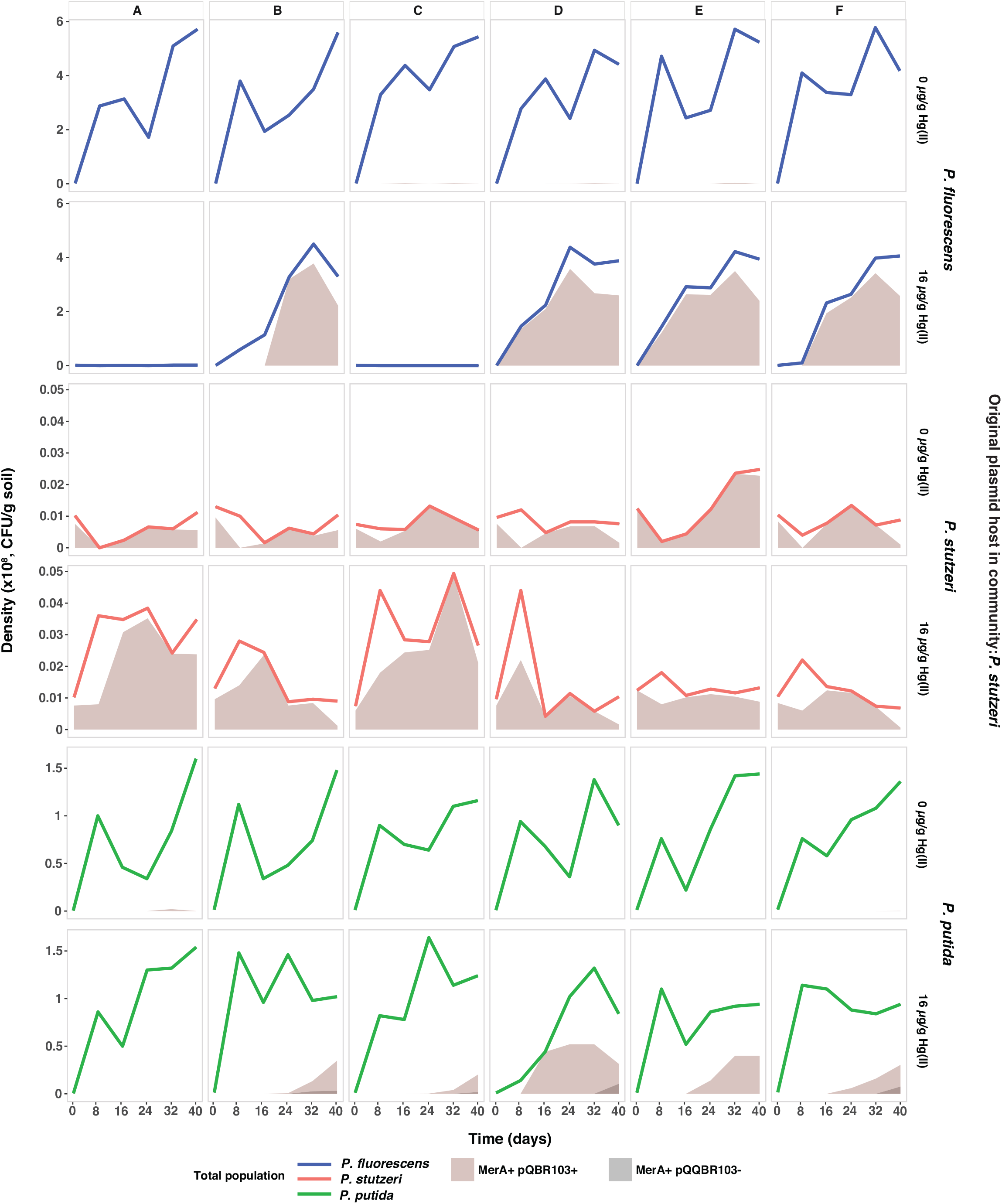
Population density and mobile genetic element dynamics in communities where *P. stutzeri* was the original plasmid host. A-F clonal populations evolving with or without mercury. Lines show the population densities of *P. fluorescens* (blue); *P. stutzeri* (red); *P. putida* (green). Brown areas show the density of plasmid carriers; Grey areas show the density of cells that have retained the Tn5042 but lost the plasmid.

**Figure 4.**
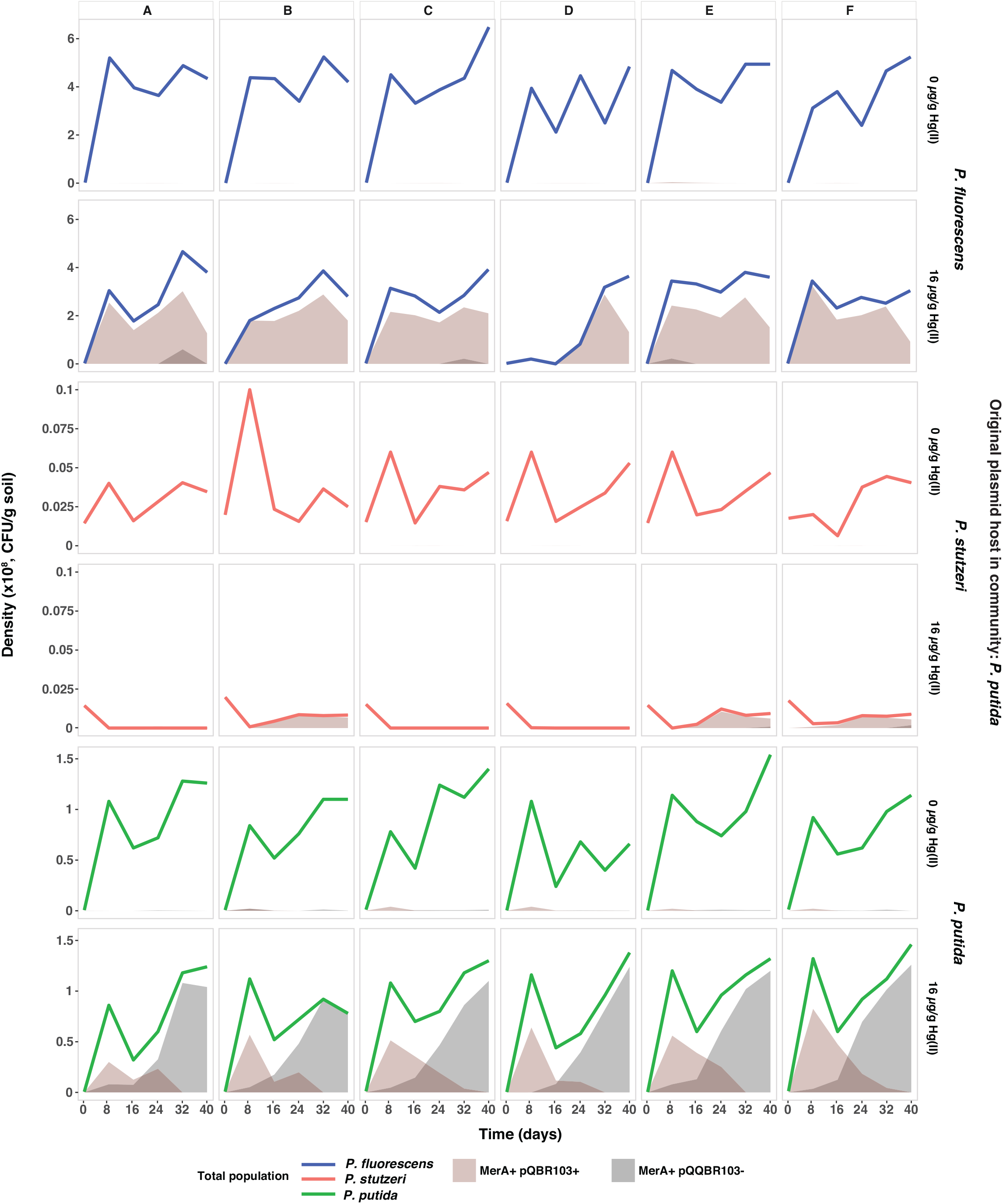
Population density and mobile genetic element dynamics in communities where *P. putida* was the original plasmid host. A-F clonal populations evolving with or without mercury. Lines show the population densities of *P. fluorescens* (blue); *P. stutzeri* (red); *P. putida* (green). Brown areas show the density of plasmid carriers; Grey areas show the density of cells that have retained the Tn5042 but lost the plasmid.

### Diversity of plasmid-carriers in communities

Finally, we tested how the original plasmid host identity affected the diversity of plasmid-carriers at the end of the experiment. The diversity of plasmid-carriers was affected by both the original plasmid host species identity (effect of plasmid treatment; 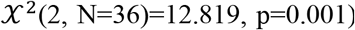 and mercury selection (effect of mercury; 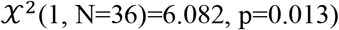 (Figure 5). Without mercury selection the diversity of plasmid-carriers was highest when *P. stutzeri* was the original plasmid host. Whereas, with mercury selection, the diversity of plasmid-carriers was higher in communities where *P. fluorescens* or *P. stutzeri* were the original plasmid hosts compared to communities where *P. putida* was the original plasmid host. Consistent with our data on community-level plasmid abundance, these data show that the diversity of plasmid-carriers is likely to be higher when plasmids are beneficial and are introduced to the community by proficient plasmid hosts.

**Figure 5.**
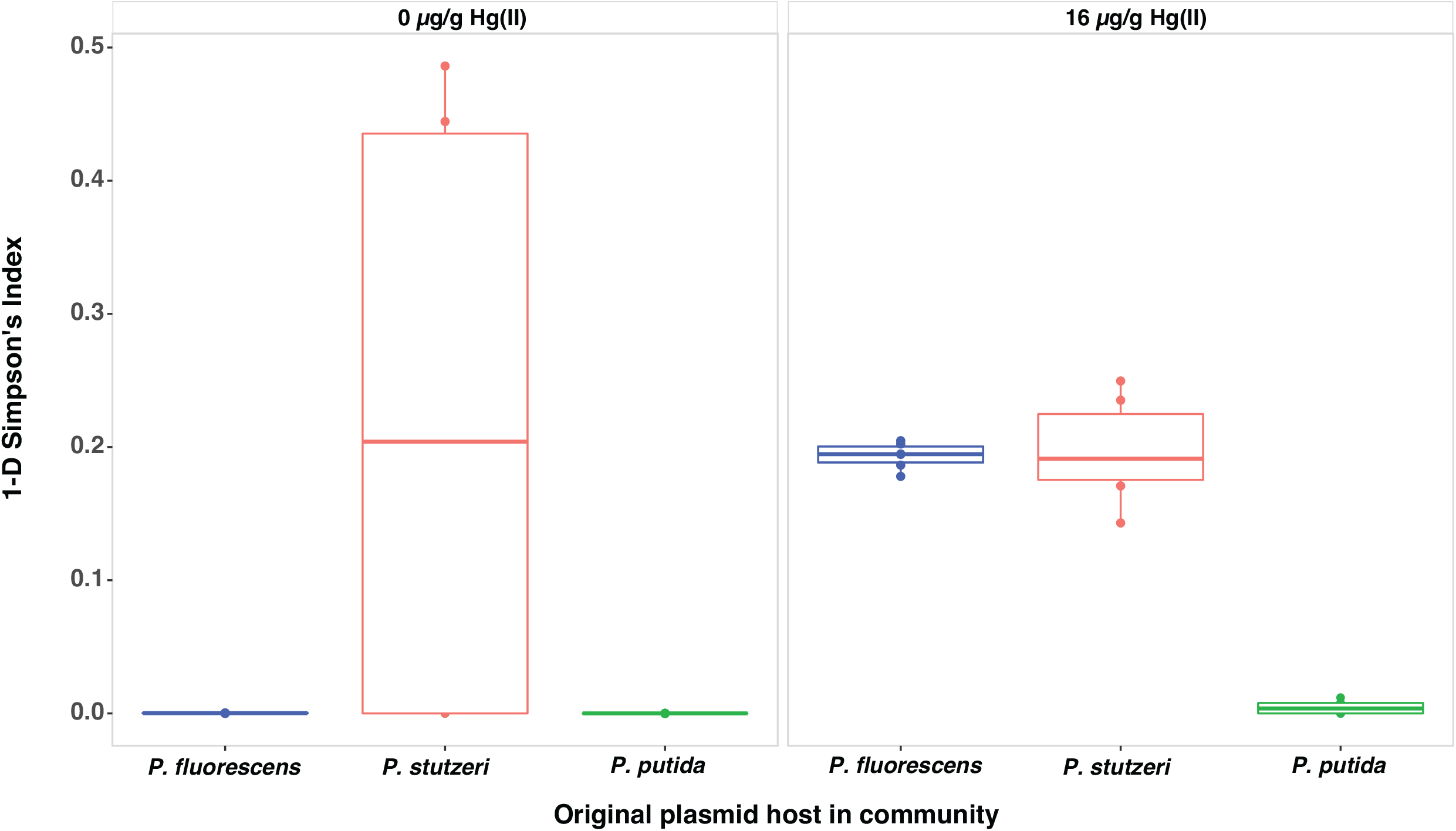
Diversity of plasmid-carriers at the end of the experiment. Species diversity was calculated as the 1-D Simpson’s Index by using the end-point population densities of the plasmid-carriers in each species in each original plasmid host community.

## DISCUSSION

In natural microbial communities, broad host range plasmids are frequently transmitted to diverse host species thus highlighting the importance of plasmids in HGT and their role in the spread of resistance genes in the environment (Klümper *et al.* 2015). In this study, we aimed to understand the extent to which plasmid dynamics in a bacterial community are affected by the original plasmid host species identity within that community. Our findings suggest that plasmid abundance at the community-level was driven by the identity of the original plasmid host species. We observed that pQBR103 reached higher community-level abundance when hosted by a proficient plasmid-host, *P. fluorescens*. Dionisio *et al.* (2002) have previously shown the importance of species identity in shaping the plasmid dynamics in a community. This was further described by Hall *et al.* (2016) where a proficient plasmid-host could act as a source of the plasmid for a non-proficient host species in a two-species soil community. These plasmid dynamics were explained in terms of conjugative plasmids persisting in the community as infectious agents via interspecies transfer (Bahl, Hansen and Sørensen 2007). Here, we extend these results to a more complex three-species community, a different plasmid, and a wider range of plasmid host species and proficiencies.

The community-level plasmid abundance also varied according to mercury selection. In common with previous studies (Cairns *et al.* 2018), plasmids were observed at higher frequencies in recipient species in the presence versus absence of positive selection. Detecting HGT events is more likely under positive selection, because, while individual conjugation events may be rare, positively selected horizontally acquired genes will rise to high frequency due to clonal expansion. This has led to a generally accepted, but probably incorrect view, that HGT is accelerated under positive selection (Aminov 2011; Fletcher 2015). By contrast, recent experimental data shows that horizontal transmission plays a more important role in plasmid stability in the absence of positive selection (Stevenson *et al.* 2017), leading to higher rates of gene mobilisation and transfer in these environments (Hall *et al.* 2017b). Mercury selection also drove the invasion of *P. putida* mutants that had lost the plasmid but captured the Tn*5042* carrying the mercury resistance operon to the chromosome, an outcome rarely observed in the other host species. This confirms our previous data that the rate and/or propensity for transposition of traits from the plasmid to the chromosome is variable among *Pseudomonas* species (Kottara *et al.* 2018). We show here that the dynamics of this process are affected by the community context, specifically whether or not *P. putida* was the original plasmid host. Chromosomal capture of mercury resistance transposon in *P. putida* occurred earlier when it began the experiment with the plasmid, reflecting that transposition is random mutational event and thus more likely to occur in larger – plasmid-bearing – populations. Interestingly, however, our data also show that even low proficiency plasmid hosts, which rapidly capture useful traits and jettison the plasmid, can act as a source of plasmids for other species in community by transferring the plasmid to more proficient host species before it is lost.

In contrast to the study of Hall *et al.* (2016), which used a highly conjugative plasmid, pQBR57, the plasmid used here, pQBR103, has >1000-fold lower conjugation rate (γ) (Log_10_(γ) pQBR103= ~ −13.8, Log_10_(γ) pQBR57= ~ −10.8; Hall *et al.* 2015). While previous studies of pQBR103 have focused on the importance compensatory evolution in its longer-term stability (Harrison *et al.* 2015), here we show an effect of between species conjugation increasing the community-level abundance of the plasmid. The role for interspecific conjugation in pQBR103 stability was most notable in communities where it was initially carried by a non-proficient original plasmid host, *P. putida*. Here, while the plasmid started in ~33% of the population and went extinct in the *P. putida* population, it survived in the community by horizontal transmission, most commonly into *P. fluorescens*. Through interspecific conjugation, pQBR103 increased the diversity of plasmid-carriers in communities, especially under mercury selection. However, this effect depended upon the original plasmid host species identity. Conjugation also depends on the population density, and in this case the higher population density of *P. fluorescens* could have enabled the plasmid transfer from *P. fluorescens*. Surprisingly, although with mercury selection more proficient plasmid host species (*i.e. P. fluorescens* and *P. stutzeri*) allowed higher diversities of plasmid-carriers, without mercury it was in communities where the moderately proficient plasmid host, *P. stutzeri*, was original plasmid host where the highest plasmid-carrier diversity was observed. This effect is likely to have been caused by the more equitable distribution of plasmid carriage in these communities, and specifically by higher rates of plasmid carriage in *P. stutzeri* itself compared to communities where this species had to obtain the plasmid via conjugation.

Soil microbial communities are highly diverse, which is thought to play a key role in their function (Torsvik and Øvreås 2002) and species diversity has been suggested to play a role in the dissemination of conjugative plasmids (Dionisio *et al.* 2002). Soil habitats are often characterised as hot spots for HGT (van Elsas and Bailey 2002; Sørensen *et al.* 2005) due to the spatially structured nature of such environments (Bahl, Hansen and Sørensen 2007; Fox *et al.* 2008; Røder *et al.* 2013). Here, we show that the identity of original plasmid host species determines the community-level abundance of conjugative plasmids in soil bacterial communities. Proficient plasmid hosts better maintain plasmids within their own population and transmit these plasmids to other species in the community. This implies that proficient plasmid host species could promote the robustness of communities by spreading potentially adaptive genes to more diverse species, allowing their survival upon environmental deterioration in the future.

## Supporting information

Data for Figures 1-4

Data for Figure 5

## ACKNOWLEDGEMENTS

The authors thank T. Daniell and D. Rozen for their comments on a previous version of this work.

## AUTHOR CONTRIBUTIONS

AK, JH and ΜB designed the study; AK performed the experiments and analysed the data; AK and MB drafted the manuscript.

